# Lost and found: re-searching and re-scoring proteomics data aids the discovery of bacterial proteins and improves proteome coverage

**DOI:** 10.1101/2019.12.18.881375

**Authors:** Patrick Willems, Igor Fijalkowski, Petra Van Damme

## Abstract

Prokaryotic genome annotation is heavily dependent on automated gene annotation pipelines that are prone to propagate errors and underestimate genome complexity. We describe an optimized proteogenomic workflow that uses ribo-seq and proteomic data of *Salmonella Typhiumurium* to identify unannotated proteins or alternative protein forms raised upon alternative translation initiation (i.e. N-terminal proteoforms). This data analysis encompasses the searching of co-fragmenting peptides and post-processing with extended peptide-to-spectrum quality features including comparison to predicted fragment ion intensities. When applying this strategy, an enhanced proteome-depth is achieved as well as greater confidence for unannotated peptide hits. We demonstrate the general applicability of our pipeline by re-analyzing public *Deinococcus radiodurans* datasets. Taken together, systematic re-analysis using available prokaryotic (proteome) datasets holds great promise to assist in experimentally-based genome annotation.

## INTRODUCTION

With the exponential rise of sequenced bacterial genomes, automated genome annotation pipelines are indispensable. Despite their utility, a growing body of evidence suggests that these methods underestimate genome complexity and pose a danger of propagating biases present in current annotations (Poptsova and Gogarten 2010; Warren et al. 2010; Wood et al. 2012). For instance, in the widely adopted NCBI prokaryotic genome annotation pipeline, protein start site annotation, sequencing errors giving rise to interrupted genes and the delineation of open reading frames (ORFs) based on homology or *ab initio* predictions remain persistent problems (Haft et al. 2018). In addition, genomic elements such as small open reading frames (sORFs) are vastly underrepresented in current genome annotations as emphasized by their systematic identification in recent reports studying translation in bacteria (Miravet-Verde et al. 2019; Sberro et al. 2019; Weaver et al. 2019). Resolving these annotation biases necessitates experimental studies aiming to delineate protein-coding regions. During genome re-annotation efforts, confident assignment of novel protein-coding regions is crucial. Such endeavors are greatly facilitated by OMICs techniques such as ribosome profiling (ribo-seq). Ribo-seq provides a genome-wide snapshot of *in vivo* translation through deep sequencing of mRNA fragments covered by the actively translated ribosomes and can hint towards unannotated translation products (Ingolia et al. 2009; Li et al. 2012). Furthermore, proteomics can provide complementary evidence of protein synthesis by searching customized protein databases such as full genome translations (Jaffe et al. 2004) or *de novo* assigned ORFs based on ribo-seq and/or sequence features (Ndah et al. 2017; Clauwaert et al. 2019; Miravet-Verde et al. 2019). Supported by complementary ribo-seq and proteomics evidence, novel ORFs, N-terminal extensions and truncations or wrongly annotated pseudogenes and their translation products have been reported in several bacteria (Ndah et al. 2017; Clauwaert et al. 2019).

In terms of peptide identification, correct discrimination of true positives from false positives is complicated due to the reasonably increased database size searched in proteogenomics (Nesvizhskii 2014). Re-scoring of peptide-to-spectrum matches (PSMs) by using machine learning tools, such as Percolator (Granholm et al. 2014), can aid in assigning correct peptide identifications. Such post-processing analysis uses several scoring features describing the quality of the PSM. For instance, search engines such as MS-GF+ deliver an extended set of PSM features (e.g. MS-GF+ score and matched fragment peak mass deviations), that can be used by Percolator (Granholm et al. 2014). In addition, the deviation of the predicted peptide retention time (RT) serves as a useful scoring feature (The et al. 2016). Next to predicting the RT for a given peptide, algorithms were recently introduced that predict the intensity of peptide fragment ions with unprecedented accuracy (Degroeve and Martens 2013; Gabriels et al. 2019; Gessulat et al. 2019; Tiwary et al. 2019). Comparing these predicted fragment ion intensities with those matched fragment ions aids in discriminating correct PSMs by machine learning (Gessulat et al. 2019; Silva et al. 2019; Tiwary et al. 2019). As such, these fragment intensity correlation features are especially useful to attain a higher confidence for novel (i.e. database unannotated) peptide identifications (Willems et al. 2017; Verbruggen et al. 2019).

While fragment intensity-based correlation metrics typically increase the number of tryptic peptide identifications by 5% at a 0.01 FDR significance (Gessulat et al. 2019; Tiwary et al. 2019), searching spectra for evidence of co-fragmented, and thus co-fragmented, peptides can deliver up to 30 to 64% additional peptide identifications (Shteynberg et al. 2015; Dorfer et al. 2018). The identification potential of chimeric spectra is of course dependent on many factors such as proteome complexity or instrument settings such as precursor isolation window and dynamic exclusion time (Dorfer et al. 2018). Interestingly, proteins solely identified by searching such chimeric spectra tend to display lower expression levels (Dorfer et al. 2018). As such, searching chimeric spectra might aid the identification of unannotated peptides typically missed in a routine data analysis workflow. Here, we describe how searching chimeric spectra with post-processing including MS^2^PIP-derived features improves the overall proteome depth and aids in identifying hypothetical and unannotated proteins. We apply our workflow to the well-characterized human bacterial pathogen *Salmonella enterica* serovar Typhimurium (*S.* Typhimurium) and validate novel protein-coding regions with complementary ribo-seq translation evidence. We further elaborate how (ribo)proteogenomics is instrumental for reannotating ORFs and the discovery of novel ORFs across bacteria.

## RESULTS

### Maximizing peptide identification using an iterative search strategy with Percolator post-processing

Proteomic shotgun analyses of the *Salmonella enterica* subsp. *enterica* serovar Typhimurium strain SL1344 (hereafter referred to as *S.* Typhimurium) proteome was performed, profiling three consecutive exponential growth stages (O.D. 0.2, 0.4, and 0.6). For complementary evidence of protein synthesis, we relied on our previously published ribosome profiling (ribo-seq) data acquired in similar growth conditions at an O.D. of 0.5 (Giess et al. 2017; Ndah et al. 2017). Striving to search the full complement of possible genomic ORFs, all theoretical ORFs with minimal length of 30 nt. and initiated from canonical ATG or near-cognate GTG and TTG start codons (the latter two codons, estimated to account for 14% of ORF start codons, (Ndah et al. 2017)) of the *S.* Typhimurium genome were *in silico* translated and used for database searching. The resulting database contains ∼320,000 ORF translations, though exhibiting a high level of redundancy as overlapping entries with in-frame translation starts are prominent and thus solely differ by their N-terminal protein sequence. To remove this redundancy, a non-redundant tryptic peptide database of 2.2 million peptides was constructed (see Methods). Of these, approximately 550,000 peptides (25%) match annotated (Ensembl) proteins, whereas 1,6 million hypothetical peptides (75%) stem from *in silico* genome translation. Hence, searching a six-frame translation results in solely a four-fold increased database size, in contrast to the manifold increased sizes if applying such rationale to eukaryotic genomes. To assign contaminant peptides, 8,042 tryptic peptides of the *in silico*-digested cRAP database (Dorfer et al. 2018) were appended to the tryptic peptide database.

A concatenated target-decoy peptide database was searched using MS-GF+Percolator (Kall et al. 2007; Granholm et al. 2014). Next to MS-GF+ features, we relied on an additional set of 11 features, referred to as the ‘auxiliary’ feature set and including the experimental RT deviation (ΔRT) to predictions (Moruz et al. 2010), the number of missed cleavages, features related to the number of matched b/y-ions, and features describing b/y-ion intensity correlation to those of MS^2^PIP-predicted spectra (Degroeve and Martens 2013; Silva et al. 2019). A full description of the in total 34 features is provided in supplemental Table S1. To assess the merit of the feature sets, PSMs were re-scored by Percolator using either the default MS-GF+Percolator features, the auxiliary features, or the combined feature set. After post-processing, spectral and peptide q-values were re-estimated separately for annotated and novel peptides based on the recalibrated Percolator scores. Importantly, this post-processing strategy was also used for chimeric peptide identification. To this end we implemented an iterative search strategy, similar to Shteynberg *et al.* (Shteynberg et al. 2015), where fragment ions of a confidently identified peptide (PSM *Q*-value ≤ 0.01) are removed and the resulting ‘cleaned’ spectrum searched in a next search round to identify potential co-fragmented peptides (see Methods).

Firstly, we assessed the performance of the different feature sets by the identification of annotated (Ensembl) peptides. Using the combined feature set results in an increase of 1.49 ± 0.14 % (mean ± SD) non-redundant peptides identified per sample in the first search compared to default MS-GF+Percolator processing, accumulating up to 545 additional unique peptides (+ 2.1 %) across all samples (Supplemental Table S2-3). Interestingly, the synergy of post-processing by this combined feature set is more prominent for the chimeric searches. Across all samples, an additional 2,430 (+14.4%) and 2,660 (+54%) non-redundant peptides were identified using the combined feature set compared to the default MS-GF+Percolator processing in the second and third search round, respectively (Supplemental Table S2-3). Hence, MS2PIP-derived scoring features are increasingly useful for discriminating co-fragmented peptides. For instance, next to MS-GF+ score, Pearson correlation to the MS^2^PIP-predicted spectrum is a useful independent scoring metric to distinguish high-quality PSMs (Figure 2B). This is further supported by the learned Percolator SVM weights, indicating Pearson correlation as a useful feature for discriminating target PSMs in the first and subsequent iterative searches (‘spec_pearson_norm’, Supplemental Figure S1). It should be noted that high spectral correlation values can be obtained for PSMs with few matched fragment ions. This issue can (partially) be compensated by including features related to the number of identified ions (Supplemental Table S1). Taken together, re-scoring PSMs using the auxiliary feature set identified 98.5% (25,493/25,894) of the PSMs identified using default MS-GF+Percolator processing (peptide *Q*-value ≤ 1%), whereas 998 (+ 5.9%) and 1,174 (+ 23.8%) additional non-redundant peptides were identified in chimeric search rounds 2 and 3, respectively (Supplemental Table S2-3).

**Figure 1.**
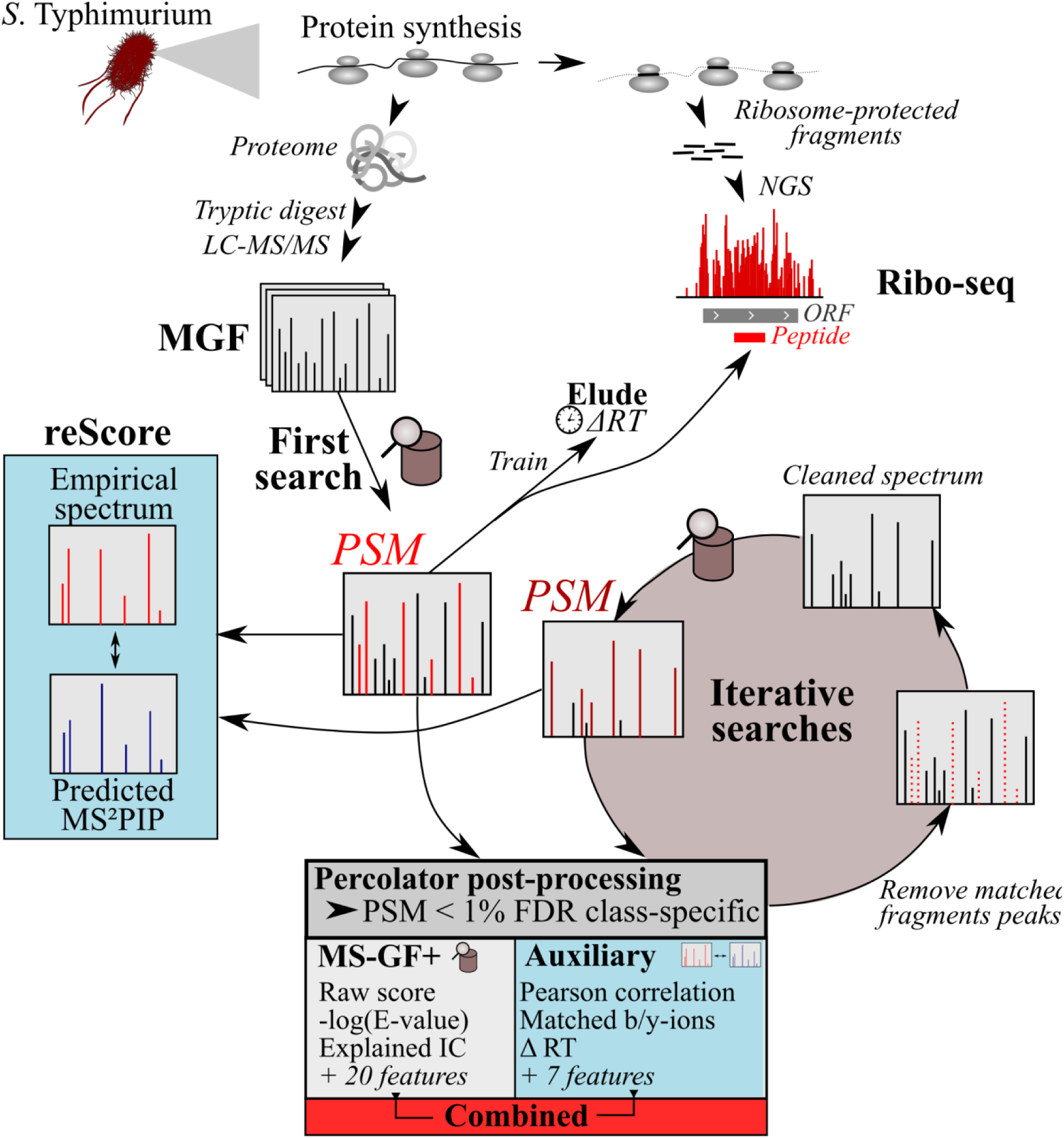
Proteomics pipeline using Percolator (Kall et al. 2007) post-processing of 34 features (supplemental Table S1). *S*. Typhimurium protein expression is studied by ribosome profiling (ribo-seq) data (Ndah et al. 2017) and proteomic shotgun analysis. Spectra were searched by MS-GF+ against an *in silico*-digested tryptic peptide database (see text and Methods). The retention time (RT) of the top-scoring 1,000 non-redundant peptides (highest MS-GF+ score) are used to train a RT model with ELUDE (Moruz et al. 2010) and calculate the deviation of empirical and predicted RT (ΔRT). Besides ΔRT, an additional ten PSM quality features were measured, constituting the ‘auxiliary’ feature set. The MS-GF+, auxiliary or the combined feature set was used by Percolator (Kall et al. 2007) for re-scoring of PSMs. *Q*-values were re-estimated in a class-specific manner for annotated and novel peptides. Identified fragment ions are removed from spectra with a significant PSM (*Q*-value < 0.01, combined feature set) and searched iteratively to identify co-fragmented peptides. Per search, identical search and post-processing steps are repeated as for the first search, except that the trained RT model is used from the first search and the application of a wider-precursor mass tolerance (similar as described by Shteynberg *et al.* (Shteynberg et al. 2015).

**Figure 2.**
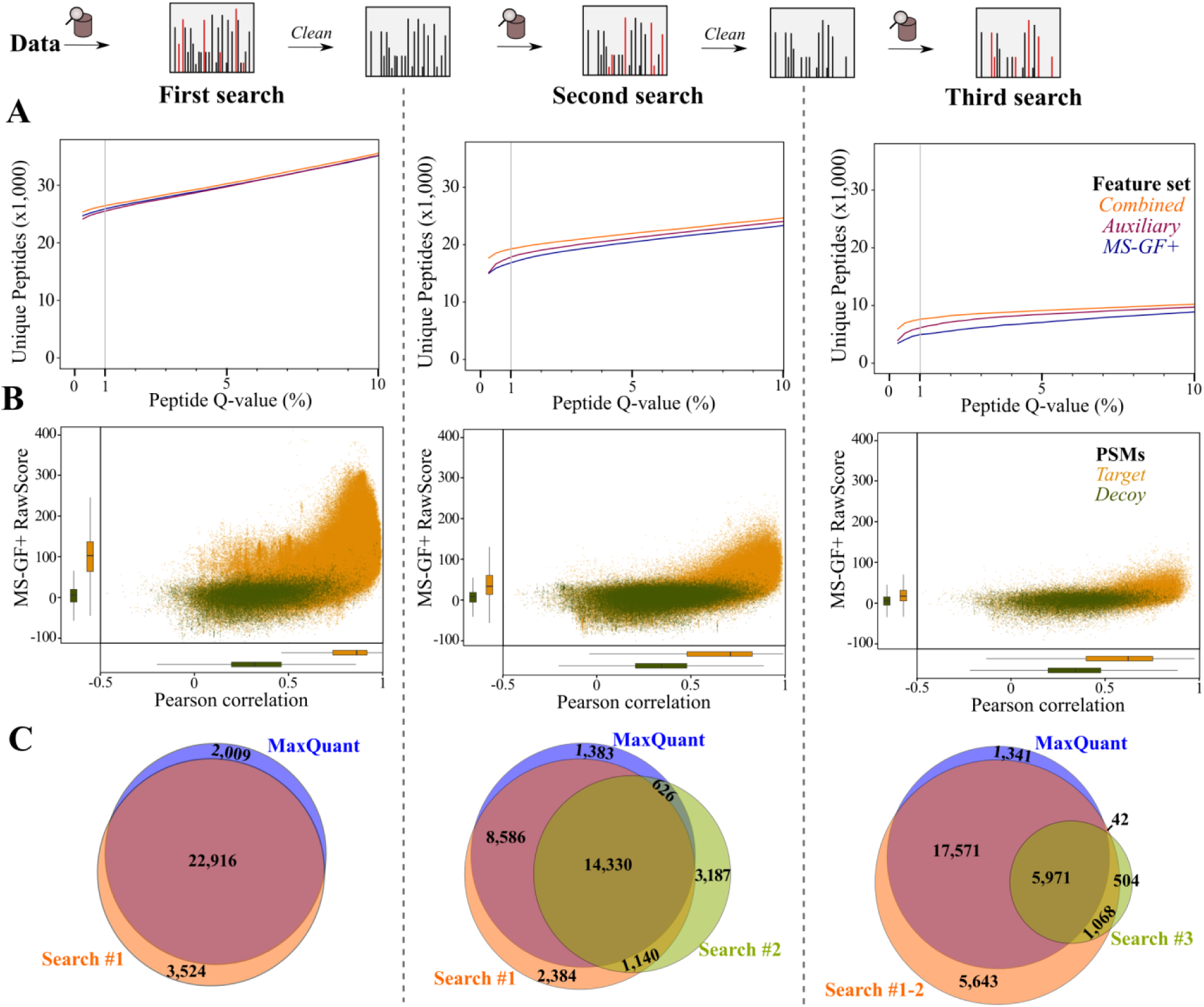
Annotated peptide identification using a chimeric post-processing pipeline. (**A**) Number of non-redundant peptide identifications (y-axis) at Percolator peptide Q-value thresholds (x-axis) in the first (left), second (middle) and third search (right). Percolator was ran in parallel using the default MS-GF+ features (blue), the auxiliary features (purple) and the combined feature set (orange). (**B**) Scatterplot of MS-GF+ RawScore and Pearson correlation (‘spec_pears_norm’ by reScore (Silva et al. 2019)) for PSMs in the three iterative search rounds. Only features for the PSM with highest Percolator re-calibrated score were displayed. (**C**) Overlap between peptides identified by MaxQuant and the three search rounds of the proteomics pipeline (combined feature set, peptide *Q*-value ≤ 0.01).

As a benchmark, we validated the performance of our pipeline by comparing identified annotated peptides with those identified by a routine MaxQuant search (Figure 2C). The first search identified 26,440 unique peptides (peptide *Q*-value ≤ 1%), sharing 22,916 peptides (91.9%) with the MaxQuant output (Figure 2C), and overall 1,550 non-redundant peptides more. Whereas the second and third search identify an additional 1,572 non-redundant peptides compared to the first search, the majority of peptides (78.6%) identified in the iterative searches were also identified in the first search. Interestingly, 626 peptides identified exclusively in the second search were also identified by MaxQuant. Important to note is that the ‘second peptide search’ option is enabled by default in MaxQuant, which performs a similar iterative search of a ‘cleaned’ spectrum for MS/MS spectra that potentially contain co-fragmented peptides (Cox et al. 2011). Running an identical MaxQuant search without this second peptide search option, shows that 678 co-fragmented peptides were identified by MaxQuant (Supplemental Table S4, Figure S2). Interestingly, whereas 157 co-fragmented peptides were already identified in our first search, 411 of them were similarly only identified by our chimeric searches (Figure S2). Taken together, our implemented iterative search strategy combined with Percolator post-processing is able to increase the number of confident peptide identifications.

### Searching of chimeric spectra improves identification rate of low-abundant proteins

Identification of co-fragmented peptides can increase the confidence of protein identifications by increasing protein coverage. For instance, 3,100 proteins were identified based on the 26,440 non-redundant peptides identified in the first search round (combined feature set, Supplemental Table S3). Of the 2,175 co-fragmented peptides exclusively identified in chimeric searches and identified in at least two samples (peptide *Q*-value ≤ 0.01), 94.9% (2,064 peptides) matched to these 3,100 protein accessions identified in the first search round. Hence, chimeric searches can identify additional peptides and improve the coverage of proteins identified in the first search round. Furthermore, the chimeric searches identify 111 additional peptides mapping to 102 proteins. Interestingly, searching chimeric spectra has previously been reported to result in the detection of previously unidentified proteins with lower mRNA-seq expression (Dorfer et al. 2018). We aimed to test this hypothesis by using the complementary ribo-seq data. Firstly, we confirmed that ribo-seq expression positively correlated with protein abundance (Pearson R = 0.43, P-value < 2.2 e-16), for 2,568 non-ambiguous proteins quantified by MaxQuant (Figure 3A, Supplemental Table S5). We indeed observed that the translation level of the 102 protein accessions exclusively identified in chimeric searches by our pipeline (≥ 2 samples) shows lower translation evidence compared to proteins identified in the first search round (Figure 3B, Supplemental Table S6). As such, chimeric searches facilitate the detection of proteins with lower abundance. For instance, *astD*, a gene encoding a metabolic enzyme involved in Arg degradation, is poorly translated (ribo-seq FPKM = 0.80) and is solely matched by the peptide ‘AGLPAGVLNLVQGGR’ in iterative search rounds. Note that this peptide and protein was also identified and quantified as low-abundant by MaxQuant (log2 protein intensity = 22.84) when enabling the second peptides search option (Supplemental Tables S4). More precisely, this peptide was identified in 8 out of 9 samples at a nearly-identical elution time (∼ 106.6 mins) with the co-fragmenting peptide ‘RVVVGLLLGEVIR’ (originating from trigger factor protein) identified in the first search round (Figure 3B). Moreover, ‘AGLPAGVLNLVQGGR’ was also identified twice in the third iterative chimeric search, after identification of ‘RVVVGLLLGEVIR’ in the first round, and even another co-fragmented peptide ‘SAEALQWDLSFR’ (ribonuclease E) in the second search round (Supplemental Figure S3). Taken together, iterative searching of MS^2^ spectra improves the protein coverage for proteins and facilitates the detection of proteins with low abundance.

**Figure 3.**
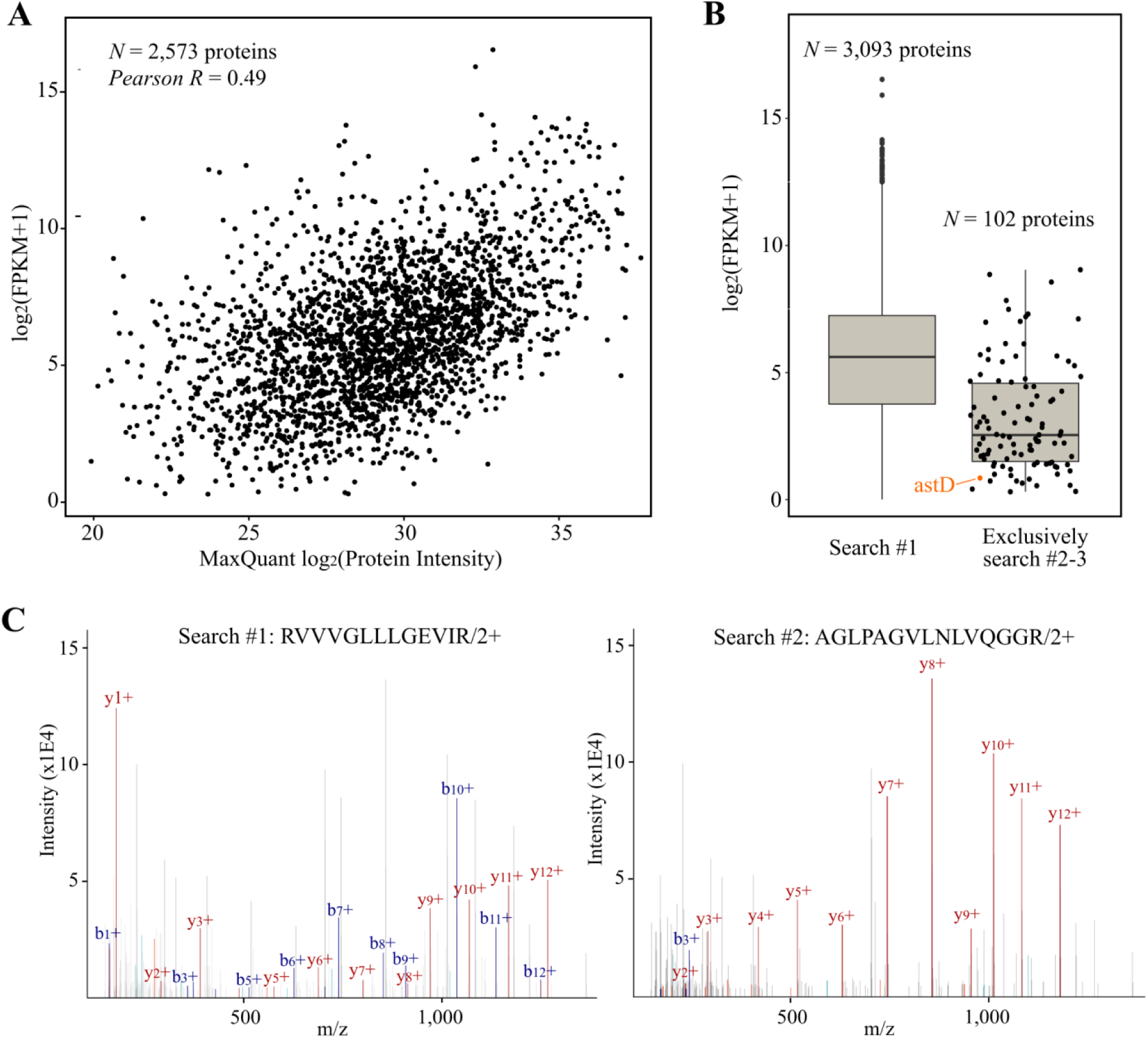
Chimeric searches improve detection of low-abundant proteins. (**A**) Pearson correlation (r =0.49) of protein abundance (MaxQuant log_2_ protein intensity, x-axis) and ribo-seq translation levels (log_2_ FPKM+1, y-axis). In total 2,573 proteins were plotted (Supplemental Table S5). (**B**) Ribo-seq translation levels for proteins matched by at least one unique peptide in the first search (including ambiguous peptide-to-protein assignments) (*Left*) and for proteins exclusively identified in the chimeric searches by at least one unique peptide in at least two samples (*Right*). The low-abundant AstD protein was indicated in orange (ribo-seq FPKM 0.80). (**C**) Annotated MS^2^ scan from the doubly charged RVVVGLLLGEVIR peptide identified in the replicate 1 sample of OD 0.8 in the first search round (*Left*) and the double charged ‘AGLPAGVLNLVQGGR’ peptide in the second search round (*Right*). Matched b/y-ions were indicated in blue and red, respectively.

### Proteogenomics

Next to searching annotated peptides, our custom peptide library comprised ∼ 1,650,000 unannotated tryptic peptides (75 % library) derived from *in silico* translation of ORFs at least 30 bp in size. Confidence of proteogenomic-indicative peptides was assessed class-specifically to conform with minimum guidelines posed for proteogenomics (Nesvizhskii 2014). Using the same post-processing strategy as described for annotated peptides, 147 and 53 peptides were identified at a 5% peptide *Q*-value by the combined feature set in the first and second search respectively (Figure 4A, Supplemental Table S7). Note that at a 5% peptide *Q*-value, no novel peptides were identified in the third search round.. Similar as the annotated peptide searches, the combined feature set improves the identification rate compared to the default MS-GF+ features (first search, +24.6% peptides and second search +60.6% peptides, Figure 4A). Whereas the MS-GF+ score and MS^2^PIP-derived Pearson correlations shows a similar distribution for target and decoy peptides, both scoring metrics aid in distinguishing high-scoring PSMs (Figure 4B). Comparing the distributions of MS-GF+ score, Pearson correlation and the fraction of explained ion current for PSMs matching annotated and novel peptides (Q-value 1% and 5%, respectively), similar distributions could be observed (Figure 4C). Hence, these PSM quality metrics indicate spectral matches to novel peptides to be of similar quality as annotated peptides.

**Figure 4.**
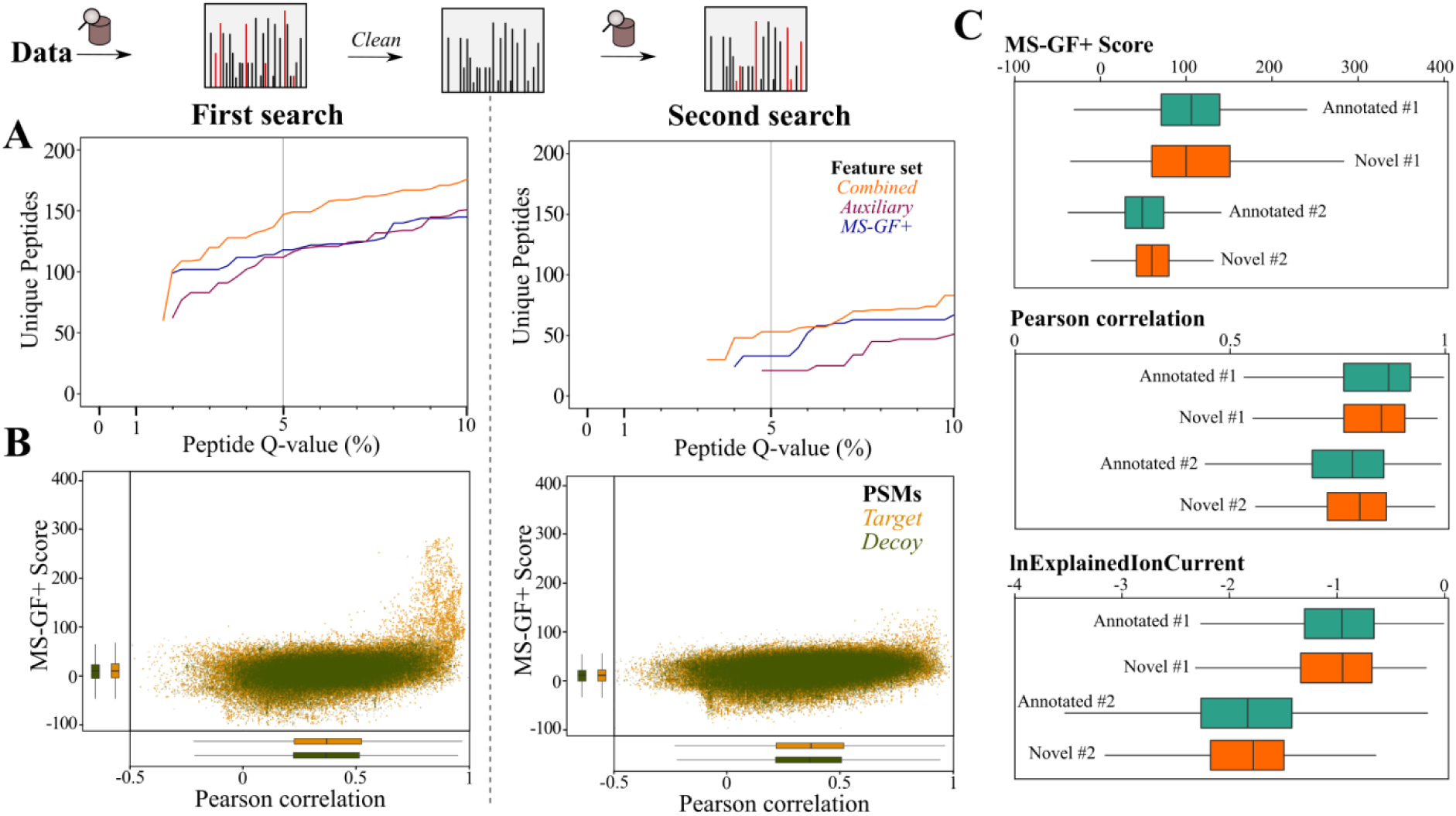
Unannotated peptide identification using a chimeric post-processing pipeline. (**A**) Number of non-redundant peptides (y-axis) at Percolator peptide Q-value thresholds (x-axis) in the first and second search. Percolator was ran in parallel using the default MS-GF+ features (blue), the auxiliary features (purple) and the combined feature set (orange). (**B**) Scatterplot of MS-GF+ RawScore and Pearson correlation (‘spec_pears_norm’ by reScore (Silva et al. 2019), supplemental Table S1) for PSMs in the two search rounds. Only features for the PSM with highest Percolator re-calibrated score were displayed after post-processing using the combined feature set. (**C**) Distributions of MS-GF+ score (*Top*), Pearson correlation and logged explained ion current (‘lnExplainedIonCurrent’) distribution for PSMs below 1% Q-value (combined feature set) for annotated peptides (green) or below 5% Q-value for unannotated peptides (orange).

In a next phase, we set out to inspect the identified peptides indicative of novel protein-coding regions in *S.* Typhimurium. To generate our final peptide candidate list, we considered the 193 unannotated peptides identified at a 5% peptide Q-value threshold using either one of the feature sets (Supplemental Table S7, Figure 5A, *Top*). In total, 942 MS^2^ spectra of representative PSMs for the 193 unannotated peptides (‘PSMIds’ in the Percolator peptide-level reports) were displayed in Supplemental Table S8. Whereas 165 out of 193 peptides (85.5%) are identified after re-scoring with the combined feature set, the MS-GF+ and auxiliary feature set deliver an additional 28 non-redundant peptides. However, the combined feature set delivers 38 extra unannotated peptides. For instance, in case of the peptide ‘SSLLSTHK’, matching a N-terminal extension of the annotated *yccA*, with an estimated peptide Q-value of 3.45% using the combined feature set, the same PSM had a peptide Q-value of 11.5% using the default MS-GF+ processing (Figure 5B). Here, the used auxiliary features, e.g. a Pearson correlation of 0.92 and a full series of identified y-ions, aided in assigning this peptide as a significant hit. Assigning peptide identifications per ORF reveals 31 unannotated ORFs matched by at least two unique peptides, 49 ORFs matched by one unique peptide with at least 2 PSMs and 48 ORFs by a peptide with a single PSM (Figure 5A, *Bottom*). To restrict our manual curation effort we ignored ORFs only matched by a single PSM, leaving 80 ORFs supported by at least two PSMs. After inspection of PSMs in the context of corresponding ribo-seq coverage and *de novo* ORF predictions (Ndah et al. 2017) in a genome viewer, we categorized 66 out of 80 novel ORFs (82.5%) as high-confident. More specifically, when classifying these ORFs with respect to Ensembl-annotated ORFs, we identified 15 novel intergenic ORFs, 1 novel ORF mapping to a so-called non-coding RNA, 38 N-terminal (Nt-)extensions, six Nt-truncations, three ORFs located at pseudogenes, two frame-shifts, and one example where a peptide over-spanned an annotated amino-acid deletion in the interrupted *yhbT* protein-coding region (Figure 5C).

**Figure 5.**
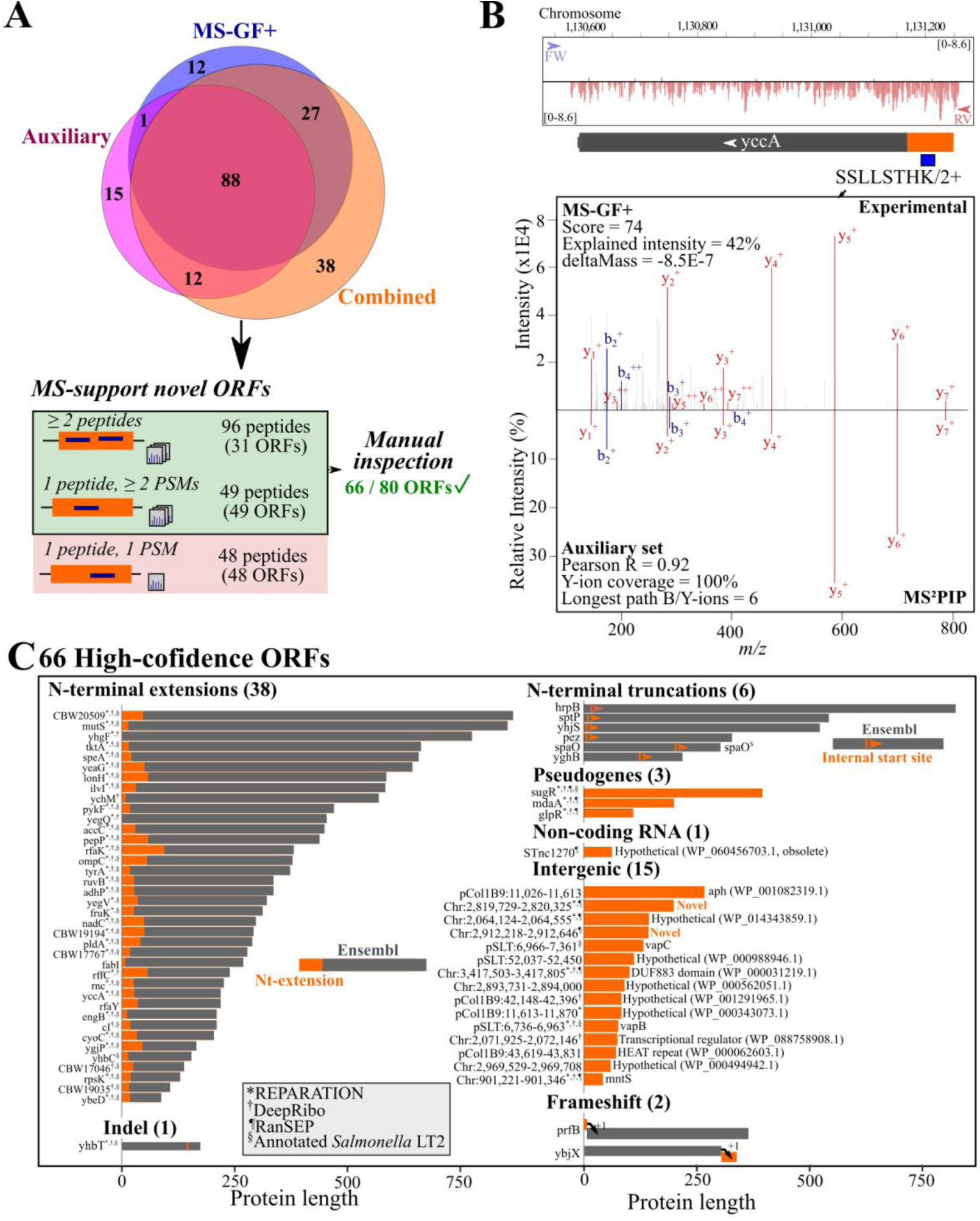
*S.* Typhimurium unannotated protein-coding regions. (**A**) *(Top*) Venn diagram of unannotated peptides identified at a peptide Q-value ≤ 0.05 after Percolator processing using the MS-GF+, auxiliary, or combined feature sets. (*Bottom*) Peptide to ORF assignment, resulting in 66 high-confidence ORFs after manual inspection (see Material and Methods). (**B**) *(Top*) Annotated MS^2^ spectrum of the doubly charged SSLLSTHK and (*Bottom*) the MS^2^PIP-predicted MS^2^ spectrum. Features used for Percolator post-processing were displayed. (**C**) Overview of 66 high-confidence unannotated protein-coding regions. Bars are indicative of protein size and grey indicates Ensembl-annotated regions, whereas orange unannotated protein regions. The corresponding Ensembl annotations were indicated on the left, whereas for intergenic ORFs, chromosomal locations and identical proteins identified by protein BLAST were displayed. In addition, was indicated whether ORF delineation corresponds to de novo predictions of *REPARATION (Ndah et al. 2017), ^†^DeepRibo (Clauwaert et al. 2019), ^¶^ranSEP-predicted ORFs (ranSEP score ≥ 0.5, (Miravet-Verde et al. 2019)) or matched S. Typhimurium str. LT2 annotation^§^.

Of the 38 Nt-extensions with matching peptide evidence, 33 (87 %) were previously predicted by REPARATION and for 14 of these, matching peptide evidence was previously reported (Ndah et al. 2017). However, we now report peptide-evidence for an additional 24 Nt-extensions with one (14) or more (10) matching peptides.. Similarly, we identified six peptides hinting towards N-terminally truncated protein variants or proteoforms. This includes for instance the functionally characterized internally translated short SpaO proteoform (SpaO^S^, (Notti et al. 2015)), which was originally described in the *Yersinia* homologue *yscQ* (Bzymek et al. 2012). In addition, there was proteomic evidence for a putative internal translation site in case of *yghB* suggesting a putative strong truncated proteoform. The other four Nt-truncations are truncations of few amino acids, of which two were predicted by REPARATION (Ndah et al. 2017). Note that unlike Nt-extensions, putative truncations are only uniquely discernible via the N-terminal peptide where a non-canonical start codon encodes for an initiator Met (and thus not Val [GTG] or Leu [TTG]).

We also controlled whether the 66 high-confident novel ORFs were annotated in the closely related *S.* Typhimurium LT2 model strain. To this end, we performed a genome alignment using Mauve (Darling et al. 2004) (Figure S4A) and inspected corresponding ORFs. Remarkably, 32 out of 38 extensions (but no truncations) were correctly annotated in strain LT2, suggesting probable misannotations in case of SL1344 (Figure 5C, indicated by §). For instance, the VapB and VapC proteins on the virulence pSLT plasmid were correctly annotated in strain LT2 (Figure S4B). Notably, a 42 amino acids-long intergenic ORF (Chromosome:901,221-901,346) is experimentally characterized as the manganese transporter protein MntS in *E. coli* (Waters et al. 2011). However, the corresponding genomic region is lacking annotation in both *S.* Typhimurium SL1344 and LT2 strains (Supplementary Figure S4C). A similar annotation conflict is the frameshift in polypeptide chain release factor 2 (*prfB*), which is experimentally characterized in *E. coli* (Craigen and Caskey 1986), but lacking annotation in *S.* Typhimurium (Figure S6A). More specifically, three peptides matched a short ORF overlapping the N-terminus of prfB in another reading frame (Figure S6A). Interestingly, the peptide ‘FRPHSNANPPGPAPAKPR’ possibly reflects an unannotated frameshift event in case of *ybjX*, reconstituting the *S.* Typhimurium redundant protein entry WP_129468947.1 (Figure S6B). It was identified in two out of three OD 0.4 replicates at an identical elution time. Due to two missed cleavages, likely due to less efficient cleavage before Pro, its peptide precursor has a 4+ charge state and several doubly charged fragment ions are observed (Figure S6B).

In case of the three ORFs located at pseudogenes, strong peptide evidence was provided before for SugR matching protein-coding *sugR* in *S.* Typhimurium LT2. Whereas, *mdaA* and *glpG* are also protein-coding in *S.* Typhimurium LT2, our ribo-seq and proteomics evidence suggests truncated forms in *S.* Typhimurium SL1344. For instance, a drastically shortened GlpR protein (110 amino acids) is translated of which the initial 107 amino acids equal the N-terminal region of the 256 amino acids-long *S.* Typhimurium LT2 GlpR protein (Figure 6A). Another interesting case was a peptide matching a ‘non-coding RNA’ STnc270 suggesting a protein that was identical to an obsolete NCBI record (WP_060456703.1). Other protein BLAST searches against the bacterial NCBI RefSeq database revealed however non-obsolete entries to be identical to 10 unannotated ORFs identified here besides VapB, VapC, and MntS (Figure S5B-C). All ten are redundant, so-called ‘multi-species’ protein entries, of which 7 hypothetical proteins were predicted by gene prediction algorithms. Hence, many of these hypothetical ORFs likely are *bona fida* protein-coding genes in *S.* Typhimurium and perhaps several other bacterial species. Interestingly, a remaining two intergenic ORFs did not correspond to any protein. However, the longest, intergenic Chr:2,819,729-2,820,325 ORF encoding an 199 amino acid long protein, shared 70% identity with a hypothetical protein in *Klebsiella pneumoniae* lacking any known functional protein domains (WP_142762344.1). Furthermore, strong complementary evidence of Chr:2,819,729-2,820,325 translation is provided for this ORF by ribo-seq and proteomics (Figure 6B). Besides a strong ribosome density matching translation of the newly identified ORF on the reverse strand of the annotated CBW18741 gene, two unannotated peptides were identified: ‘HALQILNR’ was identified in six out of nine samples and one PSM was found for the second peptide ‘SGNPQHLPMSTELFAVPEVFSK’ (Figure 6B).

**Figure 6.**
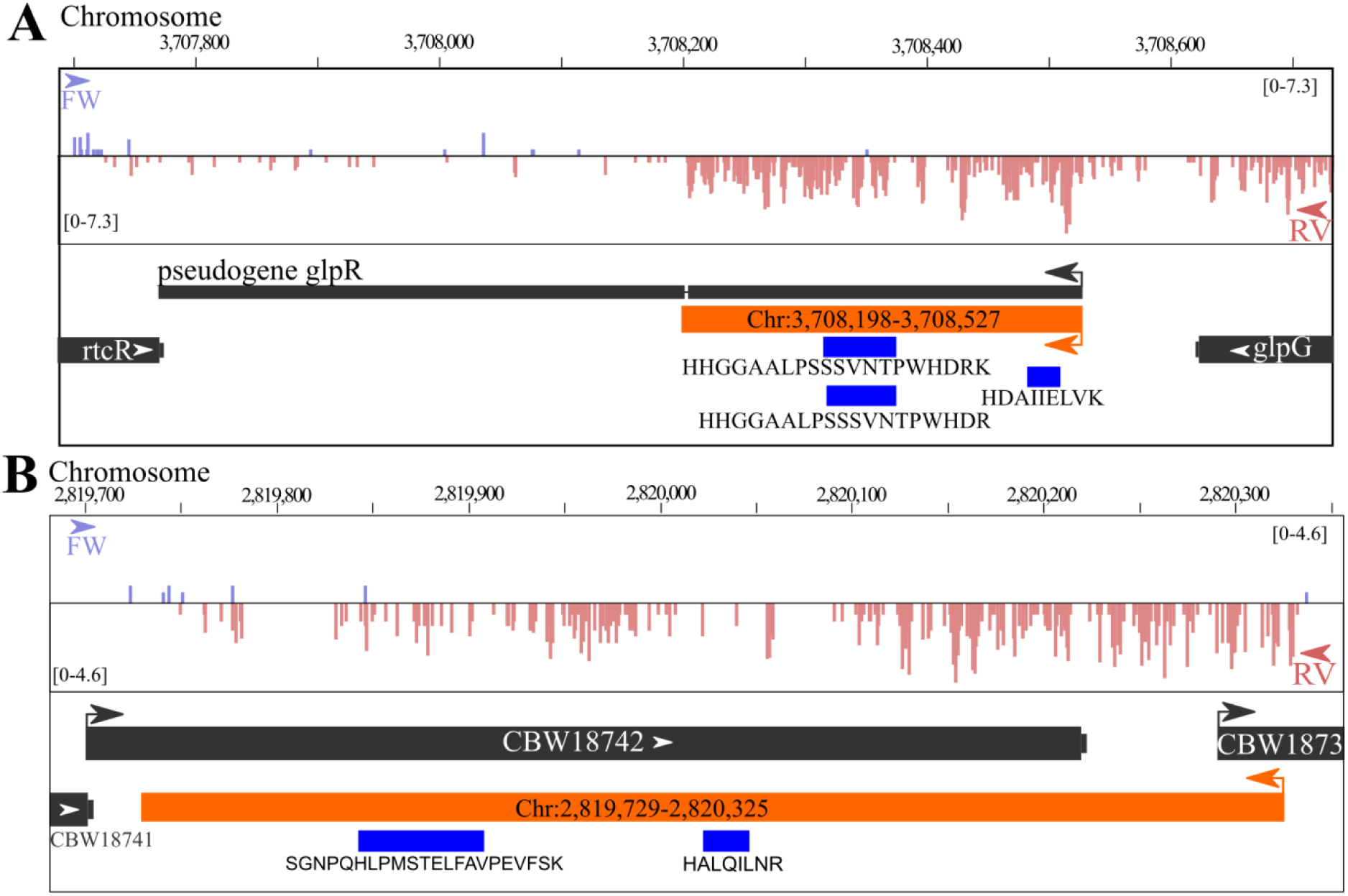
IGV (Robinson et al. 2011) genome view of ribo-seq read density and identified unannotated peptides for Chr:3,708,198-3,708,527 (A) and Chr:2,819,729-2,820,325 (B).

### Proteogenomics test case: *Deinococcus radiodurans*

Given the expanding number of sequenced bacterial genomes, a multitude of genome annotations are simply propagated. Given certain differences in annotation pipelines, e.g. different gene prediction algorithms and manual curation efforts, specific ORFs may be consistently wrongly annotated or unannotated. Proteogenomics can be a powerful tool to correctly delineate ORFs, next to the identification of novel ORFs as shown here in case of *S.* Typhimurium. Interestingly, for diverse bacterial species, a vast and expanding resource of proteomics data is accessible through public repositories for proteomics data such as PRIDE (Vizcaino et al. 2016) (Figure 7, accessed June 2019). Next to *Salmonella* (26), *E. coli* (> 100 datasets) and *Pseudomonas* species (81 studies) represent well-studied bacterial species in PRIDE. However, several other less-studied clades sometimes have high-quality shotgun datasets available and might be readily used for proteogenomic purposes. For instance, we tested the general applicability of our proteogenomic pipeline to a proteomic dataset of *Deinococcus radiodurans* str. R1 (PRIDE accession PXD011868; (Ott et al. 2019)) (Figure 7 – bold). This bacterium belongs to the *Deinoccocus-Thermus* phylum of bacteria that are highly resistant to extreme environmental conditions, though are relatively underrepresented in terms of proteomic datasets available (at least four data sets, accessed June 2019). Despite this, extremophile bacteria such as *Deinococcus radiodurans* are gaining interest for biotechnological applications due to their high resistance to UV radiation, desiccation, and oxidative conditions (Jin et al. 2019) among others. In addition, in contrast to the current *E. coli* paradigm for bacterial translation that an mRNA includes an untranslated leader sequence (UTR) harboring the ribosome-binding site (RBS) upstream of the translation initiation codon, *Deinococcus* species show a remarkably high proportion of leaderless mRNAs lacking 5’ UTRs. More specifically, up to 1,174 leaderless transcripts (encoding 60% of genes) were expressed in *Deinoccoccus desertii* (de Groot et al. 2014), whereas bioinformatics analysis suggested more than 40% leaderless mRNAs in *Deinoccocus radiodurans* (Zheng et al. 2011). As such proteogenomics efforts in *Deinococcus* may inform on alternative modes of bacterial translation.

**Figure 7.**
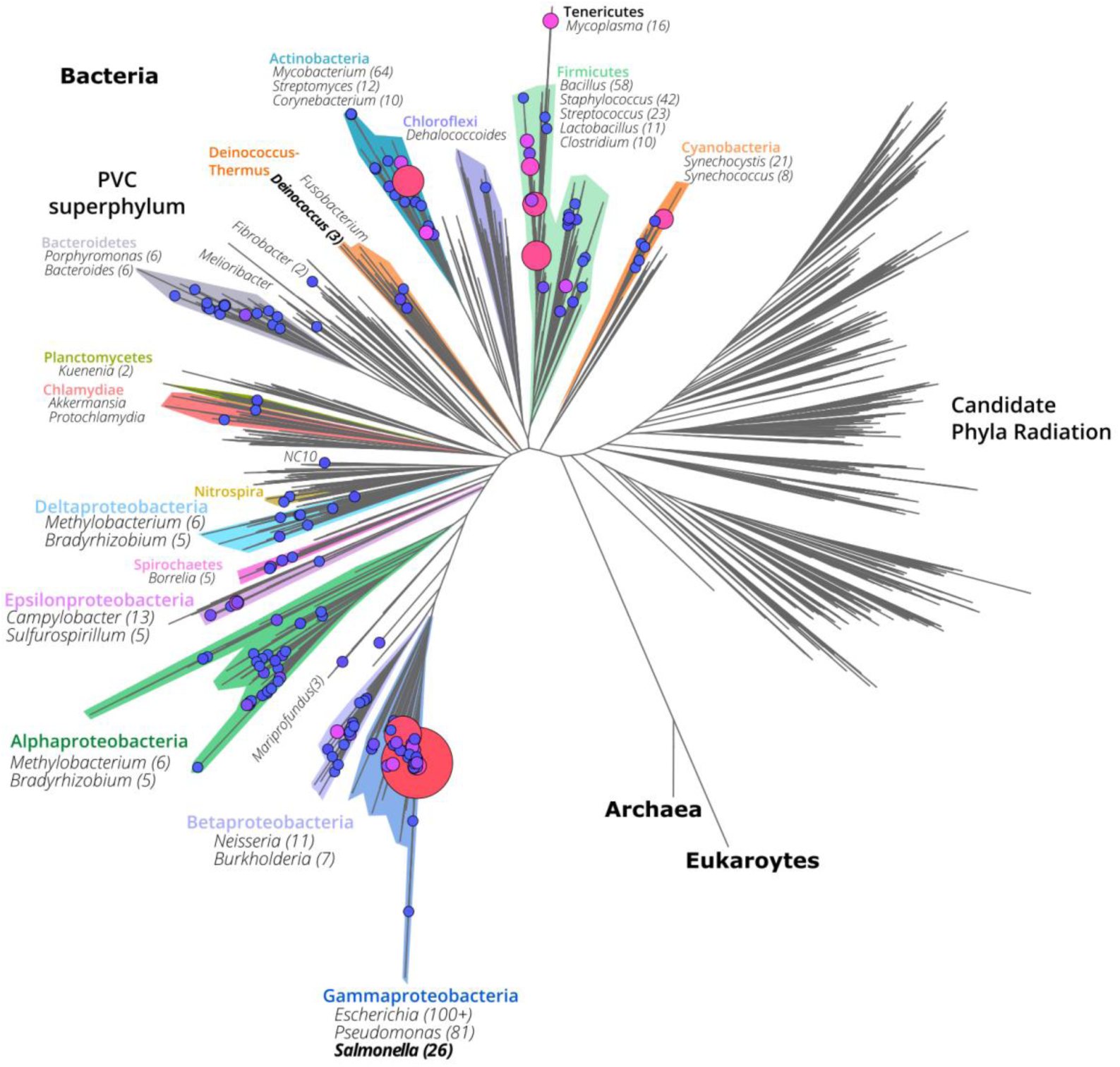
Proteomic shotgun datasets across the bacterial phylogeny available in the PRIDE repository. (Vizcaino et al. 2016). The bacterial phylogeny was adapted from Hug et al. (Hug et al. 2016), omitting the Archaea and Eukarya for ease of visualization. PRIDE accessions of bacteria were retreived and catalogued using NCBI taxonomy identifiers. Proteomic dataset identifiers plotted are given in Supplemental Table S9 with node sizes corresponding to the number of available datasets and plotted using FigTree (version 1.4.3. (2009), http://tree.bio.ed.ac.uk/software/figtree/).

Here, we re-analyzed a proteomic dataset studying the response to simulated vacuum conditions (Ott et al. 2019). As additional meta-data, we used a recent mRNA-seq dataset studying the effect of hydrogen peroxide (Lin et al. 2016). *Deinococcus radiodurans* str. R1 contains two chromosomes, a megaplasmid and a small plasmid, totaling 3,284,156 bp (White et al. 1999) (a 35% reduction compared to *S.* Typhimurium SL1344 genome size). Applying our described proteogenomic pipeline on a 6-FT database as described for *S.* Typhimurium (Figure 1, Methods), we identified 1,071 Ensembl-unannotated peptides at a 5% peptide *Q*-value using the combined feature set, of which 101 solely identified as co-fragmenting peptides in search rounds 2 or 3 (Supplemental Table S10). Notably, 273 out of 1,071 (25.5%) were annotated in the NCBI RefSeq proteome annotation (ASM856v1) of this strain, but lacking in the Ensembl annotation. Assigning these 798 NCBI and Ensembl-unannotated peptides to the longest *in silico* translated ORF yields 59 putative novel ORFs supported by at least two peptides (Supplemental Table S10). These 59 putative ORFs match nine Nt-extensions (15%, Figure 8A), a smaller proportion compared to our proteogenomic results obtained for *S.* Typhimurium (38/66 [58%] unannotated ORFs). Thanks to the high occurrence of leaderless mRNAs, the mRNA-seq coverage can provide direct complementary evidence of probable translation start sites in certain cases. For instance, in case of the annotated ORF DR_1392, translation (and transcription) starting from an upstream canonical start codon is supported by two N-terminal peptides (with and without initiator methionine (iMet) processing) and clear evidence of leaderless mRNA-seq expression (Figure 8B). Besides, 17 putative ORFs were located in genomic regions annotated as containing frame shifts (Figure 8A). However, the unannotated ORFs identified here clearly results from the incorrect annotation of a gene interruptions due to sequencing errors. For instance, the ORF 1:2,419,332-2,419,991 had eight matching peptides, whereas also upstream novel peptides were identified in an authentic frameshift region (Figure 8C). mRNA-seq coverage clearly indicates a bp deletion (Figure 8C, zoomed region), giving rise to an interrupted CDS. In fact, this sequencing error was pointed out earlier experimentally and the correctly reconstituted DncA was characterized as an essential nuclease involved in growth and radiation resistance of *Deinoccocus radiodurans* (Das and Misra 2012). Despite this reported finding, the NCBI RefSeq NP_296138.1 CDS is incorrectly translated due to the sequencing error persisting in the original genome sequence. Finally, the remaining 33 putative ORFs were located intergenic with reference to Ensembl annotated ORFs (Figure 8A); either overlapping annotated ORFs out-of-frame (11 ORFs) or antisense (18 ORFs), or non-overlapping between annotated ORFs (4 ORFs). For instance, the 90 amino acid-long encoding ORF 1:264,981-265,250 was matched by four peptides and matching mRNA-seq expression evidence (Figure 8D). In contrast, the predicted antisense DR_0263 ORF lacked any peptide match or mRNA-seq evidence. In this case, DR_0263 could thus potentially be a false prediction, with DR_0262 as last protein-coding ORF of the sense transcript and the novel 1:264,981-265,250 ORF part of an antisense transcript following DR_0264. Moreover, the novel peptide ‘ELEDLSEAEWLR’ (Figure 8D) suggests another antisense 82 amino acid-long protein-coding ORF (1:264,758-265,003, one peptide match) on this transcript, or alternatively a frameshifted protein translated from these two neighboring ORFs. For 16 out of 18 ‘antisense’ ORFs, no supporting mRNA-seq or proteomic evidence is apparent for the annotated ORF, suggesting incorrect gene predictions. In the two other cases where annotated ORFs have matching evidence, peptide and mRNA-seq coverage suggest non-overlapping truncated forms of both ORFs to be a more likely scenario. For instance, several peptides matched the annotated *deoD*, where four novel peptides matched the putative antisense ORF 2:264,981-265,250 (Figure 8E). Closer inspection of mRNA-seq coverage suggests matching leaderless mRNA transcripts with canonical ATG start codons for the annotated *deoD* and the novel ORF more likely than the ORFs corresponding with translation being initiated at the upstream, in-frame near-cognate GTG start sites, thus finally resulting in non-overlapping protein-coding ORFs on opposite strands (Figure 8E). In case of some out-of-frame sense overlapping ORFs, peptide evidence was discovered for both ORFs, suggesting possible partial overlaps. For instance, the putative ORF 1:2,518,632-2,518,976 likely encodes a 115 amino acid-long protein from a leaderless transcript that, at the C-terminus, overlaps with DR_2516 supported by a single peptide matching the annotation (Figure 8E). However, it is not always possible to discriminate protein start sites and further (matching) experimental evidence and manual curation would be required for the comphrehensive delineation of ORFs in this species.

**Figure 8.**
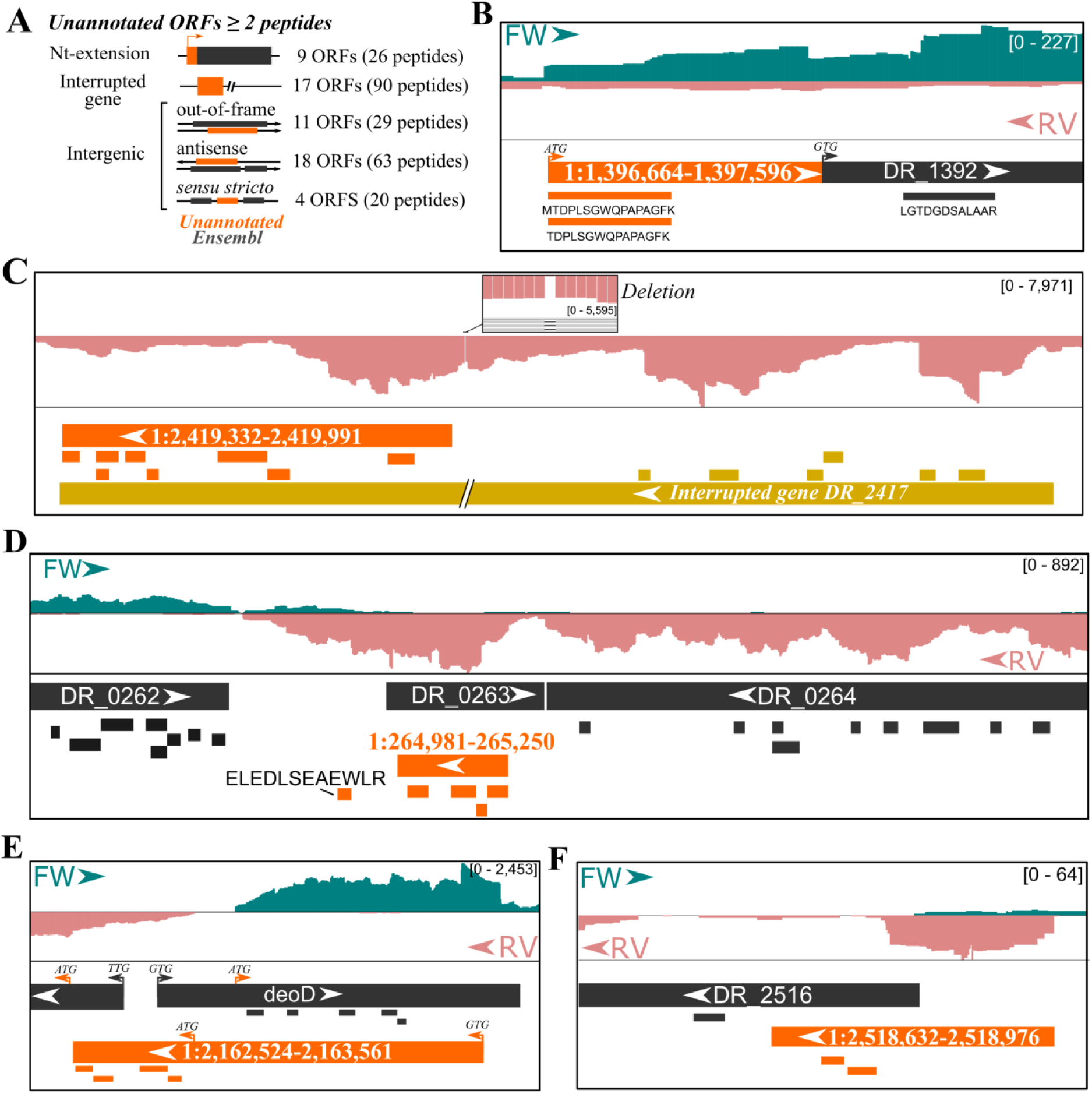
Putative ORFs in *Deinococcus radiodurans* str. R1 with matching proteogenomic evidence. (**A**) Categorization of 59 putative ORFs with at least two matching peptides not present in Ensembl or NCBI annotation (assembly ASM856v1). (**B-F**) Genome view by IGV (Robinson et al. 2011) showing identified annotated (dark grey) or novel (orange) peptides and their matching (longest) ORFs. Putative novel translation start sites were indicated by arrows with the respective start codon. In addition, stranded mRNA-seq coverage was displayed from unstressed wild-type *D. radiodurans* str. R1 (Lin et al. 2016). Respective genome region coordinates were (**B**) 1:1,396,647-1,396,869, (**C**) 1:2,419,754-2,421,036, (**D**) 1:264,080-266,689, (**E**) 1:2,162,416-2,163,711, and (**F**) 1:2,518,399-2,519,021.

## DISCUSSION

Automated prokaryotic genome annotation is indispensable given the exponential increase of sequenced bacterial genomes. Despite their utility, they can propagate inconsistent gene annotations across bacterial species (Poptsova and Gogarten 2010; Richardson and Watson 2013; Breitwieser et al. 2019; Salzberg 2019). Here, we apply an optimized proteogenomics workflow for bacteria to identify, unannotated protein-coding ORFs and correct existing annotations. Proteogenomic searches solely require a genome sequence, for which all possible canonical or near-cognate start codon initiated ORFs are searched (≥ 30 bp). Similar annotation-less searching for such small protein products has been performed before in *Mycobacterium pneumoniae* (Miravet-Verde et al. 2019). However, we perform class-specific FDR estimation of annotated and novel peptides, which is a basal guideline in and essential to achieve high sensitivity in proteogenomics searches (Nesvizhskii 2014; Li et al. 2016). Whereas a threshold of two unique peptides was posed to filter true novel ORFs (Miravet-Verde et al. 2019), high-scoring peptides nonetheless revealing true positive ORFs were not considered further. For instance, of the 49 putative ORFs in *S.* Typhimurium matched by a single peptide (with at least 2 PSMs), 35 were labeled high-confidentwhen considering additional meta-data such as matching ribo-seq data (Ndah et al. 2017), *de novo* ORF predictions (Ndah et al. 2017; Clauwaert et al. 2019; Miravet-Verde et al. 2019), protein homology of the unannotated ORF and quality of the corresponding fragmentation spectrum. Comparing the fragmentation of synthetic peptides can additionally be used to validate such one-hit wonders (Friedman et al. 2017). Due to the recent development and accuracy of peak intensity prediction tools such as MS^2^PIP (Degroeve and Martens 2013) or Prosit (Gessulat et al. 2019) - the latter trained on synthetic peptides - the comparison of predicted and empirical peptide fragmentation spectra is automated and can deliver additional PSM scoring metrics to be used by semi-automated machine learning tools such as Percolator (Gessulat et al. 2019; Silva et al. 2019). Notably, we used correlation to MS^2^PIP-predicted spectra before to discriminate high-confidence unannotated peptides in *Arabidopsis thaliana* (Willems et al. 2017). In line with our previously reported findings when making use of the PROTEOFORMER pipeline (Verbruggen et al. 2019), our proteogenomic pipeline, illustrates how Percolator rescoring increases the number and confidence of (un)annotated peptide identifications (Figure 2, 4).

Co-fragmenting peptides are widespread in MS^2^ spectra and several algorithms have been developed to mine this traditionally untapped source of peptides. In our proteogenomic effort, we used an iterative search strategy similar to Shteynberg *et al.* (Shteynberg et al. 2015) that removes fragment peaks identified in a previous search round (Figure 1). Such iterative searching identified 19,283 and 7,585 non-redundant peptides in a second and third search round when searching *S.* Typhimurium proteomics data, respectively (Figure 2). Comparison to the 678 peptides identified thanks to the second peptide search by Andromeda (Cox et al. 2011) shows 568 out of 678 co-fragmenting peptides (83.8%) to be identified by our search strategy (Figure S3). It should be noted that the second search in Andromeda is more strict as it is solely performed in case overlapping precursor peptides are detected in 3D LC-MS maps (Cox et al. 2011), whereas in our iterative search strategy all identified MS^2^ spectra (PSM Q-value 1%) are re-searched. Supporting co-fragmenting peptide identifications, 78.6% of the chimeric identifications match peptides identified in the first search round. Furthermore, 2,064 out of 2,175 peptides (94.9%) exclusively identified in iterative searches match proteins with supportive MS-evidence obtained in the first search round, which overall indicates the power of our approach to increase proteome coverage. Furthermore, the other 111 co-fragmenting peptides led to the discovery of 102 additional proteins that showed lower translation levels by ribo-seq than proteins identified in the first search round (Figure 3B). Similar results have been reported, though based on mRNA-seq expression levels (Dorfer et al. 2018). As such, identification of co-fragmenting peptides increases protein coverage as well as proteome depth and thus enables the additional identification of of proteins with low abundance. The occurrence of co-fragmented peptides is dependent on several factors such as sample complexity, employed liquid chromatography set-up and MS instrument settings. For instance, broadening the precursor *m/z* isolation width and/or shortening the dynamic exclusion time will favor the identification of co-fragmented peptides (Dorfer et al. 2018). In case of our proteomics datasets, an isolation width of 1.5 *m*/*z* and a 12 s exclusion time were used on a Q Exactive HF instrument. Although not performed here, MS instrument settings can be tuned to facilitate the identification of chimeric spectra (Dorfer et al. 2018). Another alternative to avoid the preferential selection of more abundant peptide precursors is to operate in data-independent acquisition (DIA) modus. Here, no precursor selection is performed but consecutive *m*/*z* ranges are scanned and recorded in total, reducing the discrimination of low-abundant peptides and increasing reproducibility. Also in this endeavor, MS^2^ spectrum prediction tools can prove resourceful for proteogenomics by creating spectral libraries that include potential unannotated peptides.

We first applied our proteogenomic pipeline to *S.* Typhimurium, for which we earlier reported ribo-seq assisted de *novo* ORF predictions (Ndah et al. 2017; Clauwaert et al. 2019). Our pipeline showed a drastic improvement in the proteomic detection of new protein start sites and intergenic ORFs. In our *S.* Typhimurium proteome analysis, correct protein start annotation was proven to be a major issue as we obtained evidence of 38 Nt-extended protein forms or proteoforms were corrected (Figure 5). Such correct delineation of protein start sites is a known challenge in prokaryotic annotation (Haft et al. 2018). However, comparing to the annotation of the related *S.* Typhimurium LT2 strain, 32 out of 38 Nt-extensions corresponded to annotated LT2 ORFs. Likely, the misannotation of protein N-termini in case of SL1344 originates from historical reasons, as the genome assembly of S. Typhimurium LT2 used an updated gene annotation strategy including the reassessment of start predictions (McClelland et al. 2001). Notably, the updated annotation of the SL1344 by PGAP (version 4.4, updated October 2018) resolved these 32 protein start sites, emphasizing the need to compare and select the most optimal genome assembly. Next to protein start sites, we also identified several novel ORFs. These ORFs encode hypothetical proteins predicted by PGAP, but also proteins experimentally characterized before in *S.* Typhimurium or related species. For instance, the small protein MntS (Figure S5C) described in *E. coli* (Waters et al. 2011) or the VapB and VapC proteins discovered on the virulence plasmid of *S.* Typhimurium Dublin (Pullinger and Lax 1992) and later experimentally verified in *S.* Typhimurium (Winther and Gerdes 2009). Notably, the small size (42 amino acids) of *mntS* is likely a determining factor hindering its automated gene annotation, as small ORFs are under-represented in prokaryotic genomes (Miravet-Verde et al. 2019). However, also a 199 amino acid-long ORF Chr:2,819,729-2,820,325 could be identified, likely missed due to its location antisense to annotated ORFs for which no translation evidence was found in our sampled conditions (Figure 6*B*). Other unannotated ORFs well-supported by experimental evidence include alternative protein forms or proteoforms. For instance, the identified frameshift of *prfB* (Figure S6A) is a pioneering example of an efficient frameshift in *E. coli* (Craigen and Caskey 1986) and an evolutionary study showed that for ∼70% of 86 analyzed bacterial species such frameshifting is conserved (Baranov et al. 2002). Next to frameshifting, alternative translation initiation sites downstream of the canonical start codon can give rise to Nt-truncated proteoforms. For instance, identification the well-characterized truncated form of SpaO (Bzymek et al. 2012; Notti et al. 2015) was supported by an N-terminal peptide matching translation at the internal translation site, despite not being annotated in the LT2 and SL1344 genome annotations thus far. During inspection of putative ORFs, ribo-seq serves as a useful complementary evidence track as illustrated for *mntS* (Supplemental Figure S5C), the truncated *glpR* gene (Figure 6A), the intergenic antisense ORF Chr:2,819,729-2,820,325 (Figure 6B) and a multitude of other examples. Recent developments in studying translation initiation in bacteria will further facilitate precise N-terminal protein start site delineation, and concomitantly the identification of (alternative) Nt-proteoforms (Meydan et al. 2019). Taken together, with the recently discovered omnipresence of bacterial sORF translation, the translation of multiple proteoforms per gene and alternative translational decoding events such as frameshifting are largely neglected by current annotation pipelines and concomitantly, annotation databases are not properly equipped to handle these cases.

Given the ever-increasing wealth of accessible mRNA-seq, ribo-seq and proteomics data, experimental data can be adopted to improve the confidence of protein and gene annotation. In an attempt to create a representative snapshot of the bacterial proteomic datasets captured in PRIDE (Vizcaino et al. 2016), we plotted 924 datasets (accessed June 2018, Supplemental Table S8) across the bacterial kingdom phylogeny (Figure 7). We demonstrate the potential of proteogenomics in improving annotation of bacterial species based on publicly available datasets for *Deinococcus radiodurans* (Lin et al. 2016; Ott et al. 2019). This extremophile is part of the *Thermo-Deinococcus* phylum, which is, despite its interesting application potentials, relatively under-represented in terms of available proteomic datasets (Figure 7). We identified 59 high-confident ORFs with at least two matching peptides. However, very likely, many ORFs with a single peptide hit are true positives, as for instance ‘ELEDLSEAEWLR’ suggesting a 82 amino acid part of an antisense transcript (Figure 8D). Interestingly, the strong tendency for leaderless transcripts proved useful as we observed strong mRNA-seq coverage starting at N-termini of putative ORFs we identified (Figure 8B, E-F). In fact, this leaderless expression has previously been utilized to pinpoint unannotated start sites and to identify (small) proteins in *Deinococcus desertii* (de Groot et al. 2014). We also observed several, erroneous gene interruptions due to sequencing errors, such as in case of *dncA* (Figure 8C), which was in fact experimentally corrected (Das and Misra 2012). Taken together, systematic (re-)analysis of proteomic and high-throughput sequencing datasets can provide strong annotation potential. Focusing these efforts on less-characterized species or phyla will be extremely useful to strengthen reference model species that could extrapolate annotations to close relatives.

## METHODS

### Peptide database construction

Six-frame translation (6-FT) of the *S*. Typhimurium SL1344 genome (Ensembl ASM21085v2) was performed, storing all ORFs ≥ 30 basepairs initiated from ‘ATG’, ‘GTG’ and ‘TTG’ (encoding the presumed initiator Met (iMet)). The resulting 324,010 ORFs were *in silico* digested by the ‘generate-peptides’ function using the Crux toolkit (v3.2) (Park et al. 2008) using Trypsin/P specificity with two missed cleavages and N-terminal methionine excision enabled. In addition, peptide length was set from 7 to 50 amino acids and peptide mass between 500 and 5,000 *m*/*z*. Decoy peptides were generated by random shuffling of peptide sequences while maintaining the C-terminal amino acid. In total, this delivered 2,170,335 target and 2,169,158 non-overlapping decoy peptide sequences. Target peptides, and their respective decoy peptides, matching proteins of the *S.* Typhimurium Ensembl proteome (ASM21085v2, 4,672 proteins) were labeled as annotated peptides. For comparison of database size, the same digestion rules were applied to the latest human Ensembl proteome (GRCh38 annotation).

### Post-processing proteogenomics pipeline

#### First round search and post-processing

Thermo RAW files were converted to peak lists using ThermoRawFileParser (bioRxiv 10.1101/622852). Resulting MGF (Mascot generic format) files were searched against the constructed peptide database using MS-GF+ (v2019.04.18) with enzymatic cleavage disabled. Methionine oxidation was set as fixed and the top 3 PSMs were considered per scan. Using the ‘msgf2pin’ function provided by Percolator-converter package, a Percolator (Kall et al. 2007) tab-delimited input file was extracted from the MS-GF+ mzid result files, delivering in total 22 features per PSM (supplemental Table S1). For each sample, the retention time (RT) of the top 1,000 ranked unique peptide sequences (highest MS-GF+ score) was used to train a peptide RT model by ELUDE (v3.02.1) (Moruz et al. 2010). This model was used to predict the RT for all matched peptides and the absolute deviation of the predicted and experimental RT (in mins) was used as additional scoring feature in the auxiliary feature set (supplemental Table S1). In addition, the reScore algorithm (Silva et al. 2019) was used to compare the fragment peak intensities of empirical spectra with those predicted MS^2^PIP (algorithm v20190312, HCD trained model v20190107) (Degroeve and Martens 2013). Of the resulting features, ‘spec_pearson_norm’, ‘dot_prod_norm’ and ‘spec_mse’ were used in the auxiliary feature set given their relatively high weights in Percolator learning. The auxiliary feature set was further expanded with six features, including the number of Arg/Lys residues (i.e. reflecting trypsin missed cleavages) in the peptide, as well as features reflecting the number of matched b/y-ions (supplemental Table S1). Using the MS-GF+, auxiliary and combined feature sets, Percolator (v3.02.1) (Kall et al. 2007) was used to re-score PSMs for every MS-GF+ search. The feature weights of the combined feature set were displayed in Supplemental Figure S1. Subsequently, the obtained Percolator recalibrated PSM scores were used for class-specific confidence estimation for annotated (Ensembl) and unannotated peptides. To this end Percolator was ran without any learning iterations (“--max-iterations 0”) and initial weight assigned to the recalibrated Percolator score. Resulting peptide identifications were filtered at a 1% peptide *Q*-value for annotated peptides and 5% peptide *Q*-value for unannotated peptides.

#### Iterative searches for identification of co-fragmented peptides

To identify co-fragmented peptides in a MS^2^ spectrum we used an iterative search strategy resembling the rationale followed in the reSpect algorithm (Shteynberg et al. 2015). For MS^2^ spectra with an assigned PSM (Q-value ≤ 0.01) in the prior search, matching (tolerance ≤ 0.02 *m*/*z*) b/y-ions were omitted from the spectrum (including double charged fragment ions and/or ions with neutral losses). Spectra were re-searched for charge states 2+ and 3+ using MS-GF+ with parameters as described above, except setting a wider precursor mass tolerance of 3.1 Da. Such wider tolerance allows identification of non-monoisotopic peptides in the isolation window (Shteynberg et al. 2015). PSM feature generation and Percolator post-processing was performed as described above, except that the peptide RT model trained in the first search by ELUDE was used.

### Public data re-processing

Thermo RAW files from the PRIDE accession PXD011868, a *Deinococcus radiodurans* vacuum stress response dataset (Ott et al. 2019), were downloaded and processed as described above for *S.* Typhimurium. In addition, we downloaded raw sequencing data from the Sequence Read Archive (SRA) study SRP057959, studying the transcriptomic response of *D. radiodurans* to hydrogen peroxide [41]. Reads were aligned to the same Ensembl genome and reference assembly (ASM856v1) as used for the proteogenomics analysis using STAR version 2.7.3a (Dobin et al. 2013) allowing 2 mismatches. Read pairs matching to forward strand were filtered based on flags 99 and 147 and flags 83 and 163 for the reverse strand (-f option, samtools).

## DATA ACCESS

The mass spectrometry proteomics data have been deposited to the ProteomeXchange Consortium via the PRIDE (Vizcaino et al. 2016) partner repository with the dataset identifier PXD016377 (username, password: sTdgODcP).

## ACKNOWLEDGEMENTS

P.V.D. acknowledges funding from the European Research Council (ERC) under the European Union’s Horizon 2020 research and innovation program (PROPHECY grant agreement No 803972). We also thank Steven Verbruggen from the Lab of Bioinformatics and Computational Genomics (BioBix) at Ghent University (Ghent, Belgium) for help with running REPARATION.

## DISCLOSURE DECLARATION

The authors declare that they have no conflicts of interest with the contents of this article.

